# Cytosolic and ER-associated ribosomes share rRNA 2′-O-methylation landscapes across human cell types

**DOI:** 10.64898/2026.03.26.714425

**Authors:** Ülkü Uzun, Anders H. Lund

## Abstract

Ribosome heterogeneity arising from variable rRNA 2′O-methylation (2′O-Me) has been proposed as a potential mechanism for translational specialization, but whether such heterogeneity contributes to compartment-specific translation remains unknown. Here, we systematically compare the 2′O-Me landscapes of cytosolic and endoplasmic reticulum (ER)–associated ribosomes across three human cell types: HEK293 cells, H9-derived neural progenitor cells (NPCs), and neurons differentiated from these NPCs. Using detergent-based fractionation combined with RiboMeth-seq, we generate site-resolved rRNA methylation profiles for each compartment. Within each cell type, cytosolic and ER-associated ribosomes display highly similar 2′O-Me patterns, with only modest compartment-specific differences observed at 18S:462 in NPCs and 28S:2043 in neurons. Across all samples, differences in 2′O-Me patterns are more pronounced between cell types than between compartments. Together, these findings indicate that 2′O-Me does not establish a broad ER-specific methylation signature and is unlikely to be a major determinant of ribosome localization or function at the ER.

## Introduction

Ribosomes, long viewed as uniform molecular machines, are now recognized as dynamic regulators of translation with the capacity for functional specialization(Aspden et al. 2025). Such ribosomal heterogeneity can arise through multiple mechanisms, including rRNA sequence variation, incorporation of ribosomal protein (RP) paralogs, altered RP stoichiometry, and extensive rRNA modifications (Genuth and Barna 2018). Among these features, 2′O-methylation (2′O-me) represents the most abundant rRNA modification in human cells. It is catalyzed by the methyltransferase fibrillarin in complex with C/D box snoRNAs (SNORDs) that provide site specificity (Gay, Lund, and Jansson 2022; Sloan et al. 2017). Notably, many 2′O-me sites are only partially modified in the ribosome pool, generating ribosome subpopulations with distinct modification states that differ across cell types, developmental contexts, and disease states (Krogh et al. 2016; Marcel et al. 2020).

A growing body of work demonstrates that variation in rRNA 2′O-me can influence translation, cell identity, and cellular behaviour. For example, methylation at 18S:C174 modulates decoding of GC- and AU-rich codons, shaping proliferation and metabolic programs; loss of 28S:U3904 biases human stem cells toward neuroectoderm by altering translation of developmental regulators; decreased methylation at 28S:Um2402 contributes to ZEB1-driven epithelial–mesenchymal transition and a claudin-low breast cancer phenotype; and reduced methylation at 18S:1447 impairs leukemia stem-cell maintenance, extending survival in mouse models (Häfner et al. 2023; Jansson et al. 2021; Morin et al. 2025; Zhou et al. 2023). Together, these findings establish rRNA methylation heterogeneity as a key layer of translational control that shapes cell-fate decisions. However, whether such heterogeneity also contributes to compartment-specific translation within cells remains an open question.

Given the potential for compositional variation among ribosomes, an important question is how molecular heterogeneity integrates with the spatial organization of translation in eukaryotic cells (Bourke, Schwarz, and Schuman 2023). Protein synthesis is compartmentalized within distinct subcellular domains to coordinate local physiological demands, such as those occurring near mitochondria (Luo et al. 2025), the nuclear envelope (Yang et al. 2025), or synaptic terminals (Holt, Martin, and Schuman 2019). Spatial regulation ensures precise timing and dosage of protein synthesis, supports proper folding and post-translational modification, and safeguards proteostasis by limiting the accumulation of mistargeted or aggregation-prone intermediates. A major site of spatially organized translation is the endoplasmic reticulum (ER). Although ER-associated ribosomes can serve as a global translation platform, proteins destined for secretion, membrane insertion, or organellar localization are predominantly synthesized on ER-bound ribosomes (Chen et al. 2011). Ribosome association with the ER can occur through at least three non-exclusive mechanisms: (i) cotranslational targeting mediated by signal peptides or transmembrane domains recognized by the signal recognition particle (SRP), (ii) persistent ribosome–ER interactions enabling re-initiation cycles on membrane-bound ribosomes, and (iii) mRNA-dependent localization to the ER, independent of SRP or ongoing translation, potentially through interactions with RNA-binding proteins or sequence elements within untranslated regions (Sánchez, Driessen, and Wilson 2025). Together, these pathways imply that ribosomes with distinct molecular features, including specific rRNA modifications, may be preferentially recruited to or stabilized at the ER, leading to compositional differences between ER-associated and cytosolic ribosome subpopulations.

Accordingly, cytosolic and ER-associated ribosomes coexist as partially distinct translational populations with well-documented functional differences. The ER provides a specialized environment for cotranslational protein folding, N-glycosylation, and disulfide-bond formation, while minimizing aggregation of hydrophobic proteins and reducing cytosolic stress (Braakman and Hebert 2013). Translation on the ER has been reported to enhance protein yield compared with cytosolic translation in certain mammalian systems, underscoring its importance for the synthesis of long or highly structured proteins (Horste et al. 2023; Reid and Nicchitta 2012; Stephens and Nicchitta 2008). Previous work has primarily focused on how rRNA 2′O-methylation heterogeneity shapes translation across cell types, developmental stages, and disease states, but it remains unknown whether this heterogeneity is spatially partitioned across subcellular translation sites.

Here, we asked whether cytosolic and ER-associated ribosomes differ in their 2′-O-methylation landscapes, and whether such differences might contribute to compartment-specific translation. To address this, we profiled 2′-O-methylation patterns of ribosomes isolated from cytosolic and ER-enriched fractions across three human cell types—HEK293 cells, neural progenitor cells (NPCs), and neurons. Using detergent-based fractionation combined with RiboMeth-seq, we obtained high-quality, site-resolved methylation profiles from each compartment. This dataset allowed us to assess both cell-type–specific methylation heterogeneity and potential compartment-specific differences (Figure 1).

**Figure 1.**
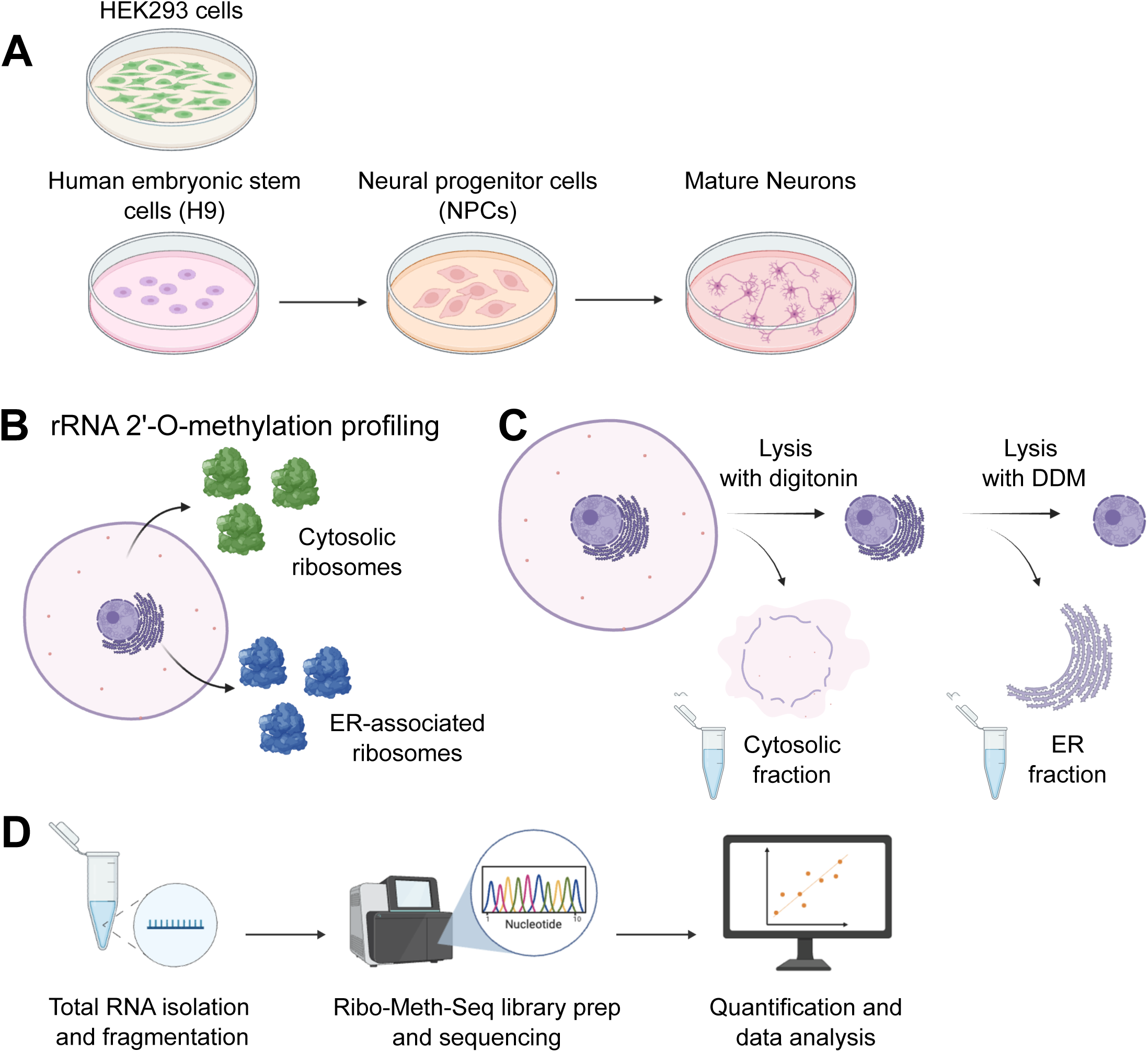
Experimental overview of cell systems, subcellular fractionation, and rRNA 2′O-methylation profiling. (A) Schematic of the cellular models used in this study: HEK293 cells as well as human embryonic stem cells (H9) differentiated into neural progenitor cells (NPCs) and mature neurons. (B) Overview of the experimental comparison between cytosolic and ER-associated ribosomes for rRNA 2′O-methylation profiling. (C) Detergent-based subcellular fractionation strategy used to isolate cytosolic and ER compartments. Cells are first permeabilized with digitonin to release cytosolic contents, yielding the cytosolic fraction, followed by solubilization of ER membranes with n-dodecyl-β-D-maltoside (DDM) to obtain the ER fraction while leaving nuclei intact. (D) Workflow for RiboMeth-seq analysis of rRNA 2′O-methylation: total RNA isolation from cytosolic and ER fractions, RNA fragmentation, RiboMeth-seq library preparation and sequencing, and subsequent quantification and computational analysis of site-specific 2′O-methylation levels.

Our results reveal that although rRNA 2′-O-methylation patterns differ markedly between cell types, ER-associated and cytosolic ribosomes within the same cell type share highly similar methylation profiles, with only subtle compartment-specific differences in NPCs and neurons.

## Results

### Isolation of cytosolic and ER-associated ribosomes

To test whether ER-associated ribosomes differ in their 2′O-methylation landscapes from cytosolic ribosomes, we used a detergent-based subcellular fractionation protocol to isolate cytosolic and ER compartments (Figure 1B,C). Cells were first permeabilized with 0.015% digitonin to release cytosolic contents while preserving ER membranes, followed by a mild 0.004% digitonin wash to minimize cross-contamination. The ER fraction was then solubilized with 2% n-dodecyl-β-D-maltoside (DDM), leaving nuclei intact (Child et al. 2021). rRNA purified from cytosolic and ER fractions was subjected to RiboMeth-seq to quantify site-specific 2′O-methylation across ribosomal RNAs (Birkedal et al. 2015). Across all three cell types, the ER fraction consistently accounted for ∼22% of total A260 absorbance, indicating a reproducible and substantial contribution of ER-associated rRNA to the total ribosome pool (Figure S1). RiboMeth-seq profiles were highly reproducible across biological replicates in all conditions (Figure S2).

### ER-associated and cytosolic ribosomes share 2′O-me patterns in HEK293 cells

We initially focused on HEK293 cells, which exhibit robust ER and secretory pathway activity (Malm et al., 2022). Successful separation of cytosolic and ER compartments was verified by enrichment of the cytoplasmic marker GAPDH and the ER marker Calnexin, as well as by qPCR for cytosolic- and ER-enriched mRNAs (Figure 2A,B) (Zinnall et al. 2022). RiboMeth-seq analysis of known 2′O-methylation sites revealed no significant differences in methylation levels between rRNA from ER-associated and cytosolic ribosomes in HEK293 cells (Figure 2C,D). As expected, approximately two thirds of sites were fully methylated, whereas the remaining sites displayed fractional methylation, indicating the presence of heterogeneous ribosome subpopulations. These observations suggest that, in HEK293 cells, the same heterogeneous pool of 2′O-methylated ribosomes engages in translation both in the cytosol and at the ER.

**Figure 2.**
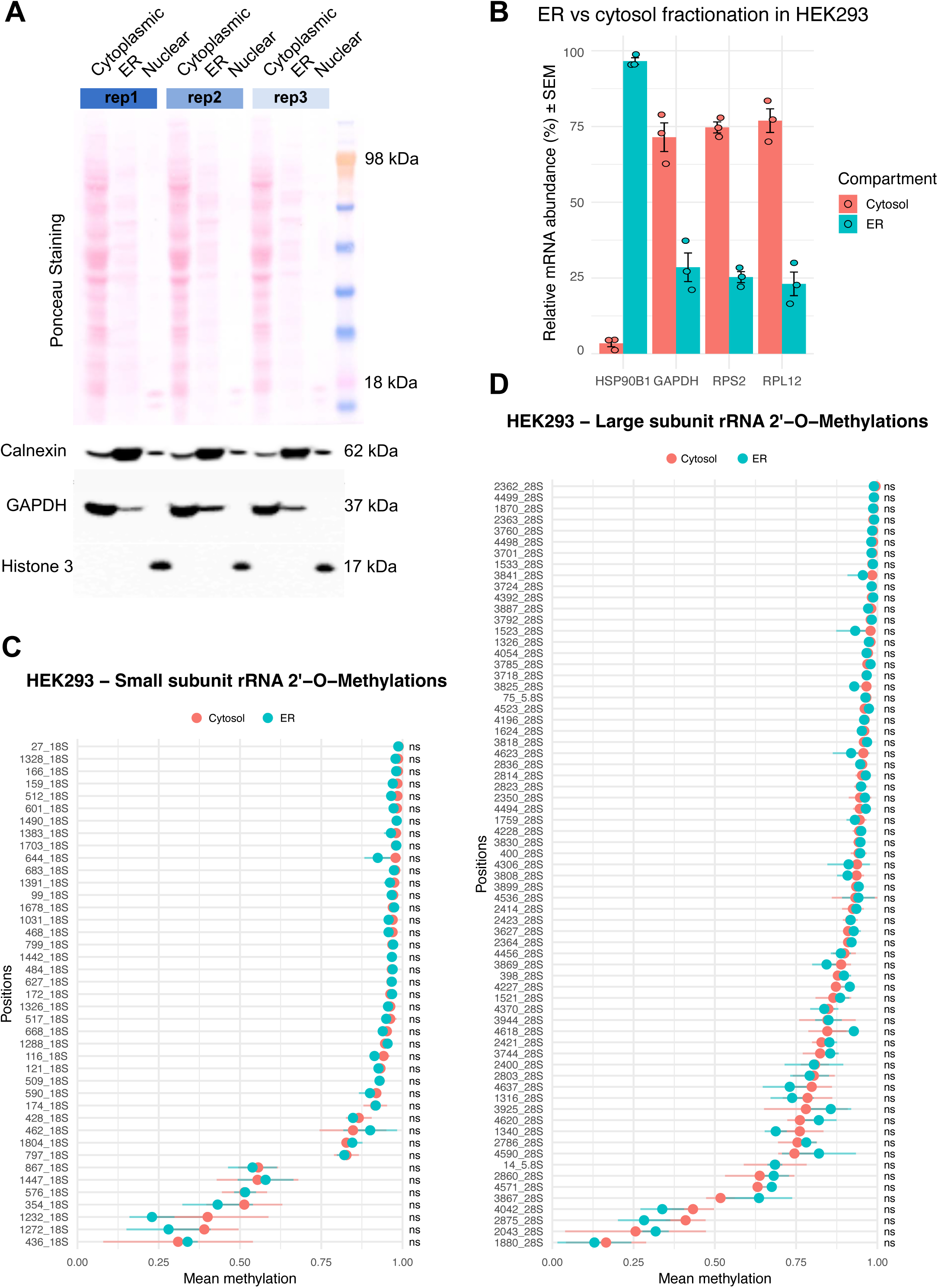
ER versus cytosolic 2′O-methylation profiles of ribosomes in HEK293 cells. (A) Detergent-based fractionation of HEK293 cells into cytoplasmic, ER, and nuclear compartments, assessed by Ponceau staining and immunoblotting for the ER marker Calnexin (62 kDa), the cytosolic marker GAPDH (37 kDa), and the nuclear marker Histone 3 (17 kDa). Three independent biological replicates (rep1–rep3) are shown. (B) Relative mRNA distribution of the ER-enriched transcript *HSP90B1* and the cytosolic transcripts *GAPDH*, *RPS2*, and *RPL12* between cytosolic and ER fractions, calculated as percentage of total signal per gene. Data represent three biological replicates. (C) Mean 2′O-methylation levels of small subunit (18S) rRNA sites in cytosolic and ER-associated ribosomes from HEK293 cells, quantified by RiboMeth-seq. Sites are ordered from highest to lowest mean methylation across all samples. (D) Mean 2′O-methylation levels of large subunit (28S and 5.8S) rRNA sites in cytosolic and ER-associated ribosomes from HEK293 cells, quantified by RiboMeth-seq. Sites are ordered from highest to lowest mean methylation across all samples. In (C) and (D), points represent the mean of three biological replicates per compartment, and error bars indicate the standard deviation. “ns” indicates no significant difference between ER and cytosolic fractions for the indicated site (two-sided t-test, p ≥ 0.05 and absolute methylation difference ≤ 0.15). 28S rRNA positions are numbered according to the human NCBI RefSeq sequence NR_003287.4 (see Materials and Methods and Supplemental Table 1 for details).

### ER-associated and cytosolic ribosomes in neural progenitor cells show largely similar 2′O-methylation profiles

We next examined NPCs, in which translational control contributes to lineage specification (Blair et al. 2017). NPCs were differentiated from H9 human embryonic stem cells, and successful differentiation was confirmed by the expression of NPC marker Pax6 (Figure 3A). Application of the fractionation protocol to NPCs again yielded well-separated cytosolic and ER fractions, as assessed by protein markers and mRNA-based qPCR assays (Figure 3B,C). In this context, RiboMeth-seq profiles of ER-associated and cytosolic ribosomes were highly similar. Among all analyzed sites, 18S:462 showed a modest but statistically significant compartment-specific difference, with methylation levels changing from 98% in cytosolic ribosomes to 83% in ER-associated ribosomes (p = 0.049), whereas no other positions exhibited significant differences in 2′O-methylation between compartments (Figure 3D,E, Sup. Figure 3). Thus, rRNA 2′O-methylation heterogeneity appears to be largely shared between ER-bound and cytosolic ribosomes in NPCs.

**Figure 3.**
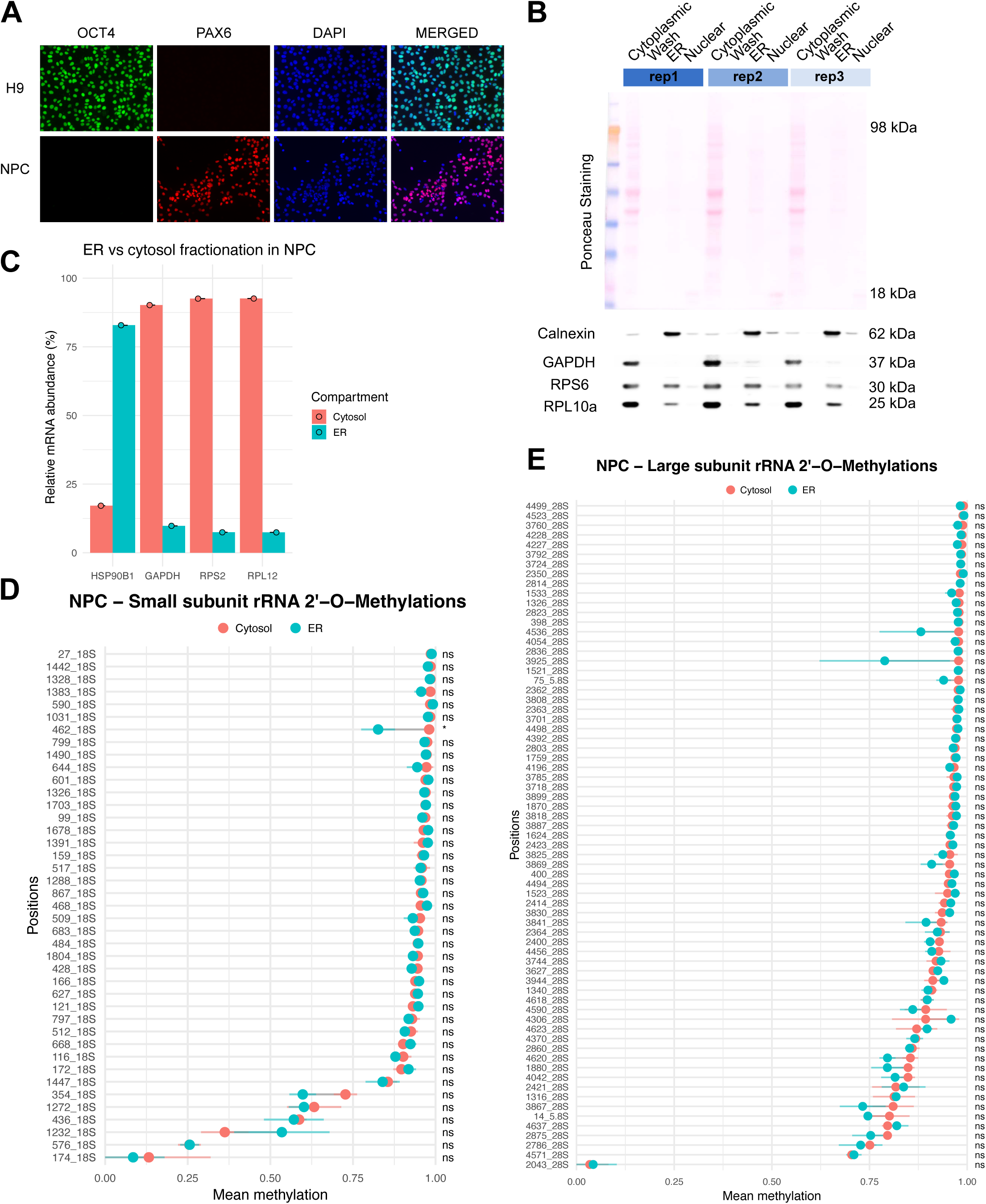
Validation of ER–cytosol fractionation and rRNA methylation profiles in human NPCs. (A) Immunofluorescence staining of pluripotent H9 cells (OCT4, DAPI) and derived neural progenitor cells (NPCs; PAX6, DAPI) confirms efficient neural induction. (B) Ponceau staining of cytoplasmic, wash, ER, and nuclear fractions from three independent NPC preparations demonstrates reproducible protein recovery across fractions, and western blot analysis of Calnexin and GAPDH verifies enrichment of ER and cytosolic markers at the expected molecular weights. RPS6, and RPL10a shows presence of small and large ribosomal proteins, respectively. (C) RT-qPCR of *HSP90B1*, *GAPDH*, *RPS2*, and *RPL12* mRNAs shows distinct relative distributions between cytosolic and ER fractions in NPCs, indicating successful compartmental separation. Data represent two technical replicates. (D) Mean 2′O-methylation levels of small subunit (18S) rRNA (E) large subunit (28S and 5.8S) rRNA sites in cytosolic and ER-associated ribosomes in NPCs, quantified by RiboMeth-seq. Sites are ordered from highest to lowest mean methylation across all samples. Points represent the mean of three biological replicates per compartment, and error bars indicate the standard deviation. “ns” indicates no significant difference between ER and cytosolic fractions for the indicated site (two-sided t-test, p ≥ 0.05 and absolute methylation difference ≤ 0.15). * indicates a significant difference (p < 0.05). For 28S rRNA numbering, see Materials and Methods and Supplemental Table 1.

### Neuronal ER-associated ribosomes display a modest 2′O-methylation shift at 28S:2043

Because neurons rely extensively on compartmentalized translation, particularly at distal processes, we next analyzed H9-derived neurons (Holt et al. 2019). Neuronal identity was validated by morphology and expression of neuronal marker Tuj1 (Figure 4A). As in the other cell types, the fractionation protocol produced distinct cytosolic and ER fractions, confirmed by marker enrichment at both the protein and mRNA levels (Figure 4B,C). Globally, RiboMeth-seq revealed that 2′O-methylation patterns were largely shared between ER-associated and cytosolic ribosomes in neurons (Figure 4D&E). Consistent with previous observations (Häfner et al. 2023), neurons generally exhibit higher overall 2′O-methylation levels (Figure 4D & E, Sup Figure 2E & F). Higher overall 2′O-methylation levels were also observed for more differentiated tissues compared to cancer cells and may be attributable to differences in the proportion of proliferating versus resting cells (Krogh et al. 2020). This could explain why maturating (non-dividing) neurons gradually accumulate higher levels of 2′O-methylation. Within this largely stable methylation landscape, a single site—28S:2043—displayed a compartment-specific difference in methylation, with 19% of ER-associated ribosomes methylated at this position compared with ∼3% of cytosolic ribosomes (p = 0.009; Figure 4E, Sup. Figure 4A). Because RiboMeth-seq scores at lowly methylated positions can be affected by 5′ and 3′ read counts at neighboring nucleotides, we examined normalized end-read profiles in a ±10-nt window around 28S:2043 and observed similar local coverage in cytosolic and ER fractions (Sup. Figure 4B), indicating that the observed difference is not driven by local read artifacts but reflects a genuine compartment-specific change in 2′O-methylation. This neuron-specific site represents a modest compartment-associated divergence within an otherwise similar 2′O-methylation profile.

**Figure 4.**
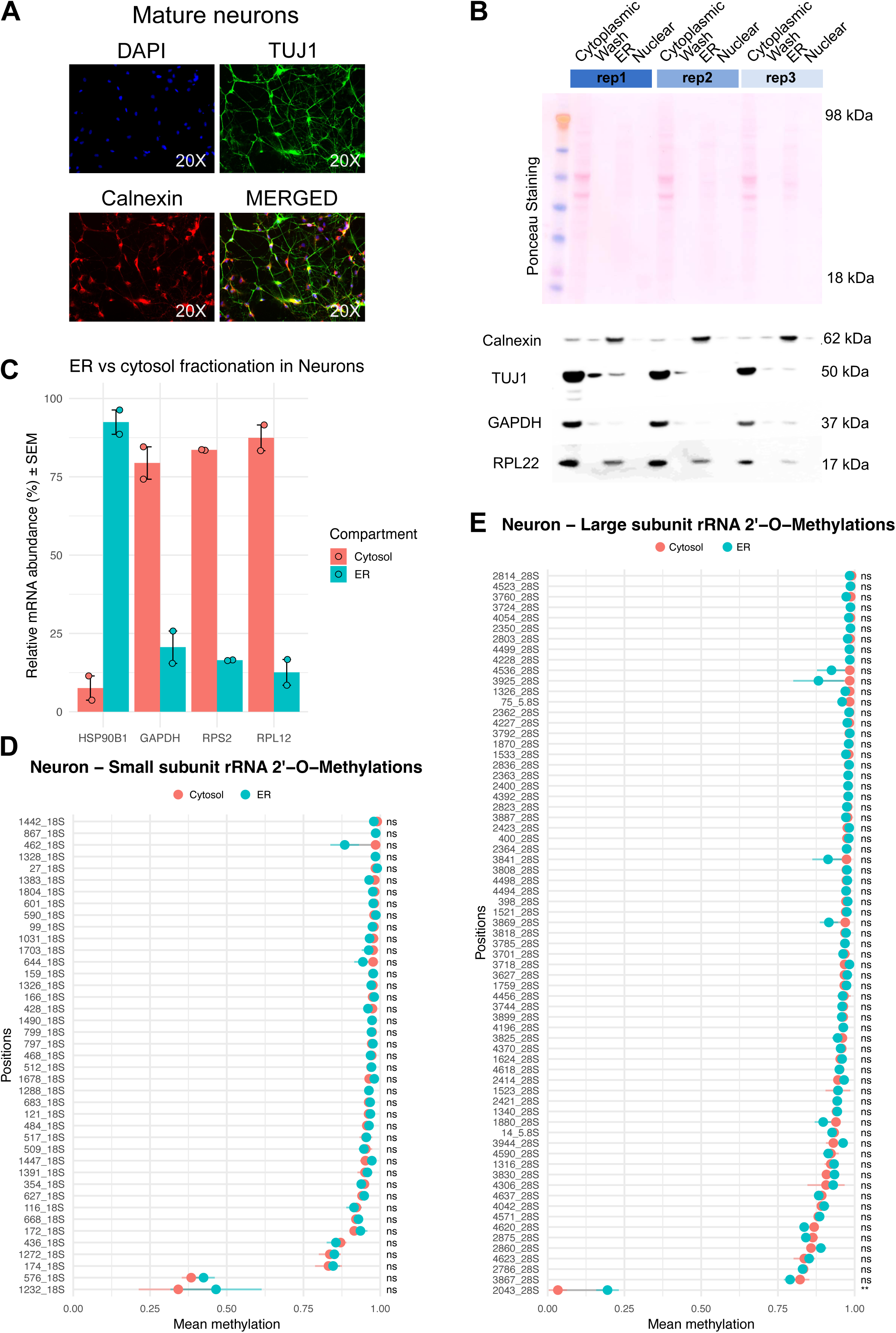
Validation of ER–cytosol fractionation and rRNA methylation profiles in mature neurons. (A) Immunofluorescence staining of mature neurons (DAPI, TUJ1, Calnexin) shows neuronal identity and ER localization in soma and neurites. (B) Detergent-based fractionation of neuronal cultures into cytoplasmic, wash, ER, and nuclear compartments, assessed by Ponceau staining and immunoblotting for Calnexin (62 kDa), TUJ1 (50 kDa), GAPDH (37 kDa), and RPL22 (17 kDa) across three biological replicates (rep1–rep3). (C) Relative mRNA distribution of the ER-enriched transcript *HSP90B1* and the cytosolic transcripts *GAPDH*, *RPS2*, and *RPL12* between cytosolic and ER fractions, calculated as percentage of total signal per gene. Data represent two biological replicates. (D) Mean 2′O-methylation levels of small subunit (18S) rRNA sites in cytosolic and ER-associated ribosomes from neurons, quantified by RiboMeth-seq. Sites are ordered from highest to lowest mean methylation across all samples. (E) Mean 2′O-methylation levels of large subunit (28S and 5.8S) rRNA sites in cytosolic and ER-associated ribosomes from neurons, quantified by RiboMeth-seq. Sites are ordered from highest to lowest mean methylation across all samples. In (D) and (E), points represent the mean of three biological replicates per compartment, and error bars indicate the standard deviation. “ns” indicates no significant difference between ER and cytosolic fractions for the indicated site (two-sided t-test, p ≥ 0.05 and absolute methylation difference ≤ 0.15). ** indicates a significant difference (p < 0.01). For 28S rRNA numbering, see Materials and Methods and Supplemental Table 1.

### Cell type and compartment patterns in rRNA 2′O-methylation

rRNA 2′-O-methylation profiles were next compared across all fractions and cell types using hierarchical clustering and principal component analysis (PCA) of RiboMeth-seq data. Hierarchical clustering visualized as heatmaps showed that samples segregate primarily by cell type, with clear separation of HEK293, NPC, and neuronal ribosomes (Figure 5A,B), indicating pronounced cell type–specific 2′-O-methylation patterns and heterogeneity. Methylation sites such as 18S:174 and 28S:1880 exhibited more dynamic changes across the different cell types. Consistent with earlier analyses, 60–70% of all methylation sites were highly methylated regardless of cell type. PCA likewise revealed clustering by cell type (Figure 5C). In contrast, ER-associated and cytosolic fractions from the same cell type clustered closely together in both the heatmaps and PCA, underscoring the overall similarity of their 2′-O-methylation landscapes. Taken together, these findings suggest that rRNA 2′-O-methylation patterns are largely shared between cytosolic and ER-associated ribosomes across different cell lines, arguing against a central role for this heterogeneity in directing ribosome localization or function at the ER, with only subtle compartment-specific changes at 18S:463 in NPCs and 28S:2043 in neurons.

**Figure 5.**
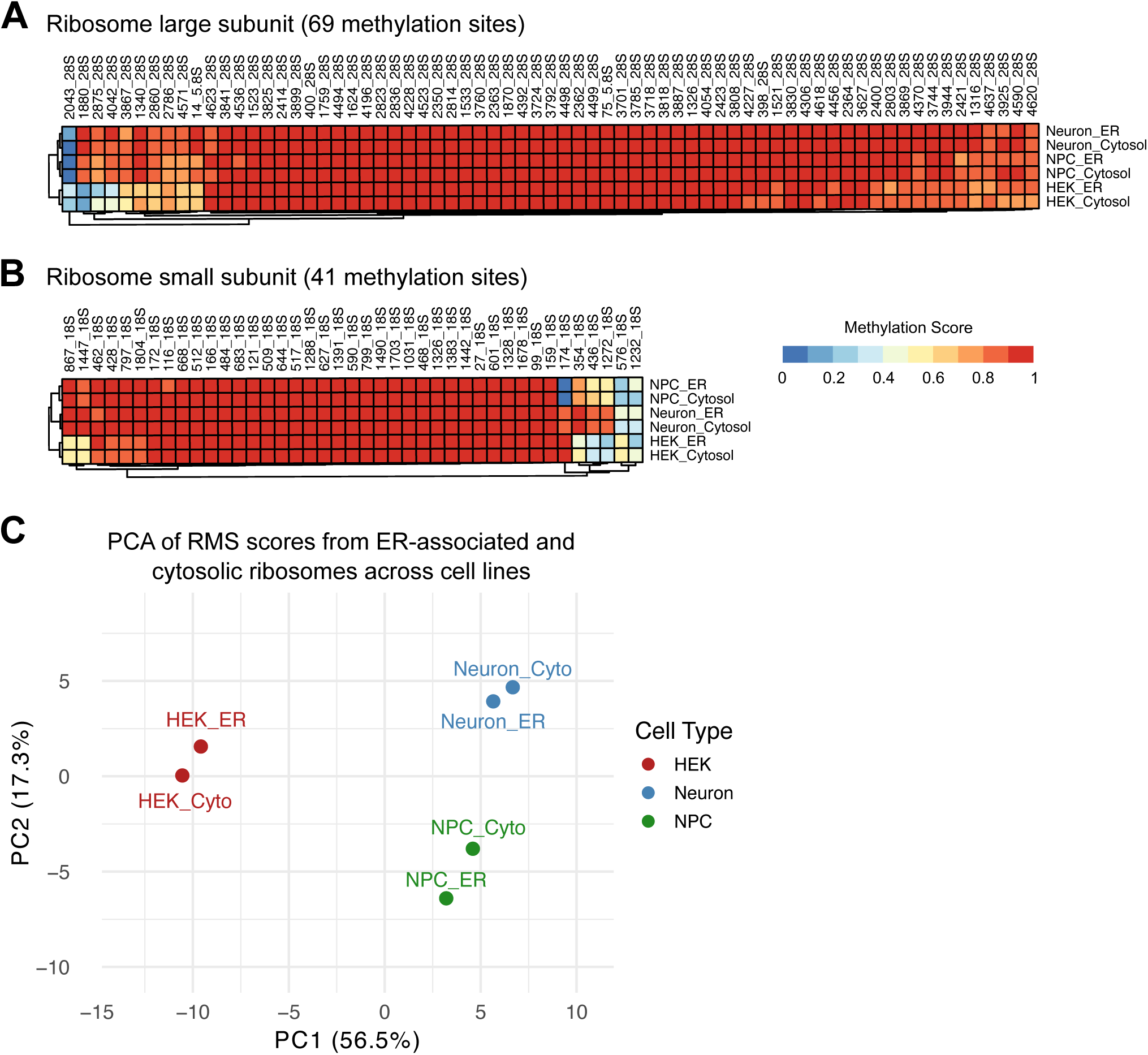
ER–associated and cytosolic ribosome rRNA 2′O-methylation profiles across cell types. (A) Heatmap of averaged signal intensities for 41 small subunit (18S) rRNA 2′O-methylation sites across six experimental conditions (HEK cytosol, HEK ER, NPC cytosol, NPC ER, neuron cytosol, neuron ER). Rows correspond to individual rRNA positions and columns to cell type and subcellular localization; for each condition, values represent the mean of three replicates, hierarchically clustered by similarity, with color indicating relative magnitude from low (blue) to high (red). (B) Heatmap of averaged signal intensities for 69 large subunit (28S and 5.8S) rRNA 2′O-methylation sites across the same six conditions, displayed and clustered as in (A). For 28S rRNA numbering, see Materials and Methods and Supplemental Table 1. (C) Principal component analysis (PCA) of RiboMeth-seq scores from ER-associated and cytosolic ribosomes in HEK293 cells, human NPCs, and neurons, showing that samples cluster primarily by cell type rather than subcellular compartment. Each point represents an individual sample; PC1 and PC2 explain 56.5% and 17.3% of the variance, respectively.

## Discussion

rRNA 2′-O-methylation has emerged as an important source of ribosome heterogeneity with the potential to modulate translation in context-specific ways. Motivated by this concept, we asked whether differences in rRNA 2′-O-methylation might contribute to spatial specialization of translation at the ER. Across the three human cell types examined, HEK293 cells, NPCs, and neurons, we did not identify a distinct 2′-O-methylation signature that defines a generic ER-associated ribosome. Instead, ER-associated and cytosolic ribosomes displayed highly similar modification profiles, with only modest site-specific differences detected in NPCs and neurons. In HEK293 cells, methylation patterns were indistinguishable between cytosolic and ER-associated ribosomes. In NPCs, a small difference was observed at 18S:462, where nearly all cytosolic ribosomes were methylated (98%) compared with 86% of ER-associated ribosomes. Although subtle, this reduction hints at a minor subpopulation of ER-associated ribosomes lacking methylation at a site located in the small ribosomal subunit, which participates in mRNA selection and decoding. Such variation could influence the translation of specific mRNA subsets within this compartment.

In neurons, compartment-specific variation was observed at 28S:2043: 19% of ER-associated ribosomes were methylated at this site, compared with only 3% of cytosolic ribosomes. Notably, this site lies within the large subunit, which engages the ER translocon (Gemmer et al. 2023). Although not directly positioned within the ribosome–ER interface, methylation at 28S:2043 may influence local structure or dynamics in ways that subtly affect ER engagement or translational regulation. Whether this modification contributes to transcript-specific translational control in neurons will require targeted functional assays.

The magnitude of these methylation differences, when considered relative to the total cellular ribosome pool, is small. Because ∼22% of ribosomes localize to the ER, a 15–16% change in methylation within this fraction represents only ∼3–4% of total ribosomes. Bulk RiboMeth-seq is limited in resolving such minor subpopulations and may partially reflect cell-to-cell heterogeneity rather than robust compartment-specific remodeling. Nevertheless, even a 3% subpopulation represents thousands of ribosomes per cell and could meaningfully impact translation of a restricted set of transcripts.

Our analysis compared cytosolic and ER-enriched ribosomes, but ribosome organization at the ER is likely far more complex. ER–ribosome interactions arise through several mechanisms—including co-translational targeting via the signal recognition particle, mRNA-dependent localization, and persistent membrane association. Each may engage distinct ribosome pools. Furthermore, structural studies have further identified at least four ER translocon complexes that interact with ribosomes depending on the class of nascent protein (Gemmer et al. 2023). Thus, rather than a simple cytosol-versus-ER methylation code, a more intricate landscape of ribosome subpopulations may exist that is masked by bulk fractionation.

Importantly, we observed partially methylated ribosomes in both compartments, supporting the presence of heterogeneous ribosome populations overall. This heterogeneity may indicate that 2′-O-methylation does not serve as a primary regulator of ER-associated translation, or alternatively, that its regulatory functions operate within smaller, highly specialized ribosome subsets that cannot be resolved with current methods.

A key limitation of our study is the reliance on detergent-based fractionation. Although this approach enriches for ER-associated ribosomes, it does not perfectly separate cytosolic, ER-bound, and other membrane-associated ribosomes (Christopher et al. 2021). Cross-contamination, even if limited, complicates interpretation of small differences in methylation. Despite this, the majority of membrane-associated translation occurs at the ER in these cells, suggesting that our ER fractions remain strongly enriched for ER-associated ribosomes.

Together, our findings argue that large-scale spatial compartmentalization of translation at the ER is unlikely to result from major differences in rRNA 2′-O-methylation. However, they do not exclude the existence of functionally distinct ribosome subpopulations within either the cytosol or the ER. Other rRNA modifications, ribosome-associated proteins, assembly intermediates, or the local biochemical environment—including tRNA availability, translation factor composition, and mRNA targeting mechanisms—may confer specificity to ER-associated translation (Reid and Nicchitta 2015). Future work integrating single-ribosome or single-cell measurements with targeted perturbation of individual modification sites will be necessary to determine how minor ribosome subpopulations contribute to ER-localized translation in both physiological and disease contexts.

## Material & Methods

### Cell lines

HEK293 cells were maintained in DMEM (Gibco, 31966-047), cells were passaged approx. 2 times a week using Trypsin-EDTA (0.25%) (Gibco, 25200056). H9 human embryonic stem cell (hESC) lines (female) were maintained on plates coated with Matrigel (Corning Life Sciences, #354277) in mTeSR Plus medium (Stem Cell Technologies,#100-0276). The culture medium was changed every other day. Cells were passaged every three days using 1X TrypLE Select (Life Technologies,#12563-011) reaching 80-90% confluence in a medium supplemented with Rock inhibitor (Y-27632) (LC laboratories, #Y-5301) at a final concentration of 10µM. For cryopreservation, cells were frozen in a solution 50% mTeSR Plus medium, 40% FBS 10% DMSO.

### Neuronal differentiation

H9 cells were diwerentiated into neural progenitor cells (NPCs) using the STEMdiw™ SMADi Neural Induction Medium (STEMCELL Technologies, #08581), following the monolayer protocol according to the manufacturer’s instructions. Briefly, H9 cells were plated on Matrigel-coated plates at a density of 2 × 10⁶ cells/cm² in neural induction medium supplemented with 10 µM ROCK inhibitor. The medium was changed daily, and the cells were passaged after 7 days. Following two additional passages (over the course of two weeks) in neural induction medium, the cells were transitioned to STEMdiw™ Neural Progenitor Medium (STEMCELL Technologies, #05833) for expansion, cryopreservation, or further diwerentiation.

H9-derived NPCs were further differentiated into forebrain neurons following the manufacturer’s instructions. First, NPCs were plated at a density of 125,000 cells/cm² on Matrigel-coated plates in STEMdiff™ Forebrain Neuron Differentiation Medium (STEMCELL Technologies, #08600) for one week with daily medium changes. The cells were then dissociated using Accutase (STEMCELL Technologies, #07920) and replated at a density of 4-6 × 10⁴ cells/cm² in STEMdiff™ Forebrain Neuron Maturation Medium (STEMCELL Technologies, #08605) on plates coated with poly-L-ornithine (Sigma-Aldrich, #P3655; 15 µg/mL in PBS) and laminin (Sigma-Aldrich, #L2020; 5 µg/mL in DMEM/F-12). The medium was changed every other day, and the neurons were matured in culture for 22 days.

### Chemical fractionation of the cytosol and ER compartments

For HEK293 cells, 7 × 10⁶ cells were seeded per 15-cm dish 2 days before harvest. NPCs were seeded at 7 × 10⁶ cells per 10-cm dish 2 days before harvest, whereas neurons were plated at 1.8 × 10⁶ cells per 10-cm dish and differentiated for 22 days before collection. On the day of collection, cells were washed once with pre-warmed growth medium (DMEM or DMEM/F12, as appropriate) and detached using Accutase for NPCs, trypsin for neurons, or by direct scraping for HEK293 cells. Cells were pelleted at 200 × g for 3 min at room temperature, resuspended in PBS lacking Ca²⁺ and Mg²⁺, and centrifuged again at 2,000 × g for 1 min at 4°C, after which supernatants were removed completely.

Lysis and wash buwers were prepared with digitonin or DDM in the common base buwer (20 mM Tris-HCl pH 7.4, 150 mM NaCl, 5 mM MgCl₂, 1 mM DTT, EDTA-free protease inhibitor cocktail [Roche, 11873580001], murine RNase inhibitor [NEB, M0314], and cycloheximide [100 µg/ml final; Sigma-Aldrich, C7698]) as described below. Digitonin stock was prepared at 2% (w/v) in DMSO (Sigma-Aldrich, D141) and stored at −20°C for up to one month, and the DDM stock (n-dodecyl-β-D-maltoside [GoldBio, DDM5]) was prepared at 20% (w/v) in cold water and protected from light and kept at 4°C.

For selective permeabilization and fractionation, pellets were first resuspended in cytoplasmic lysis buwer containing 0.015% digitonin by pipetting (20 strokes with a 200-µl pipette) and incubated on ice for 10 min. Samples were centrifuged at 2,000 × g for 5 min at 4°C, and the supernatant was collected as the cytosolic fraction, with small aliquots reserved for western blot analysis. Remaining supernatant was removed from the pellet, which was then gently washed (not resuspended) in cytoplasmic wash buwer (0.004% digitonin with the same buwer components as above) and immediately centrifuged at 2,000 × g for 5 min at 4°C. The resulting supernatant was discarded.

Pellets were subsequently further lysed with ER lysis buwer containing 2% DDM by pipetting 10 times and incubating on ice for 10 min. Samples were centrifuged at 2,400 × g for 5 min at 4°C, and the supernatant was collected as the ER-enriched membrane fraction, with small aliquots reserved for western blot analysis.

### Isolation of RNA

Total RNA isolation was performed using QIAZOL (Qiagen) and chloroform following the manufacturer’s protocol. RNA concentrations were measured using a NanoDrop One spectrophotometer (Thermo Scientific) and RNA quality were assessed using an Agilent 2100 Bioanalyzer system with the Agilent RNA 6000 Nano kit, according to the manufacturer’s instructions.

### RiboMeth-Seq

RiboMeth-seq library construction and sequencing were performed essentially as previously described (Birkedal et al. 2015; Krogh et al. 2016), except that libraries were pooled after 3′-adapter ligation containing sample-specific barcodes. Triplicate libraries were generated for each cell line or condition. A total of 1.5–4 µg of total RNA was used as input per library. RNA was partially degraded by alkaline hydrolysis at denaturing temperatures, and 20–40-nucleotide fragments were purified on 10% TBE–urea polyacrylamide gels. 3′ adapters were ligated using a system based on a modified Arabidopsis tRNA ligase that joins 2′,3′-cyclic phosphate and 5′-phosphate termini. After 3′-adapter ligation with distinct barcodes for each sample, barcoded RNA fragments were pooled, and 5′-adapter ligation was performed similarly using the modified Arabidopsis tRNA ligase. The resulting libraries were processed on the Ion Chef system and sequenced using the Ion 540™ Kit-Chef (Ion Torrent, A30011) and Ion 540™ Chip Kit (Ion Torrent, A27766).

### RiboMeth-seq Data analysis

Data were analysed as previously reported (Birkedal et al. 2015; Krogh et al. 2016). Briefly, sequencing reads were mapped to a corrected human rRNA reference sequence. The RiboMeth-seq (RMS) score represents the fraction of molecules methylated at each nucleotide position and is calculated by comparing the number of read-end counts at the queried position to those at six flanking positions on either side. Quantification in human rRNA was performed for 41 sites in 18S, 67 in 28S, and 2 in 5.8S rRNA, for which both RMS and mass-spectrometry evidence are available. Statistically significant differences in RMS signatures between two cell lines or conditions were determined by pairwise comparison (p < 0.05, two-tailed unpaired Welch’s t-test) combined with a minimum absolute change of 0.15 in RMS score. Dumbell and heatmap representations were generated in R using the ggplot2 and pheatmap packages. RMS data have been deposited in the Gene Expression Omnibus (GEO) under accession GSE325946.

The human 28S rRNA is encoded by multiple genomic copies that show sequence heterogeneity, leading to the coexistence of distinct 28S rRNA variants within a single cell. As a result, different numbering schemes have been used to annotate 2′-O-methylated residues in 28S rRNA. Earlier studies typically followed the 28S rRNA coordinates implemented in snoRNABase, whereas more recent work uses the NCBI RefSeq curated human 28S rRNA sequence NR_003287.4 (Bergeron et al. 2023), which we follow here. A correspondence table aligning snoRNABase and NR_003287.4 coordinates is provided in Supplemental Table 1; an equivalent conversion resource is also available in snoDB.

### Western blotting

Three percent of each cytoplasmic lysate, cytoplasmic wash, and ER fraction was separated by sodium dodecyl sulfate–polyacrylamide gel electrophoresis (SDS–PAGE) and transferred to PVDF membranes. The following primary antibodies were used: rabbit anti-Calnexin (Cell Signaling Technology, 2679S; 1:1000), mouse anti-GAPDH (Santa Cruz Biotechnology, sc-47724; 1:2000), rabbit anti-Histone H3 (Cell Signaling Technology, 9756; 1:1000), rabbit anti-RPS6 (Cell Signaling Technology, 2217; 1:4000), rabbit anti-RPL10A (RayBiotech, 144-05925; 1:1000), mouse anti-Tuj1 (STEMCELL Technologies, 60092; 1:500), and mouse anti-RPL22 (Santa Cruz Biotechnology, sc-136413; 1:1000).

### RT-qPCR

For validation of cytosolic and ER RNA partitioning, total RNA was isolated separately from cytosolic and ER fractions, and equal amount of RNA from each fractions was reverse-transcribed using MultiScribe Reverse Transcriptase (Invitrogen, 4311235) to generate cDNA. For RT-qPCR assays, we used the same isolated RNA pool as for RiboMeth-seq whenever sufficient material remained after library preparation, and RT-qPCR was therefore performed only on samples with spare RNA, resulting in three biological replicates for HEK293 cells, two for neurons, and one for NPCs. Relative transcript abundance was assessed by quantitative PCR (qPCR) for the cytosolic marker *GAPDH* and ribosomal proteins *RPL12* and *RPS2*, as well as the ER chaperone *HSP90B1* (for qPCR primers, see Table 1). Relative fold changes between fractions were calculated for each marker using the ΔCt method, and the percentage contribution of cytosolic versus ER fractions was derived from the normalized relative expression values, taking into account the total RNA recovered from each fraction.

**Table 1.**
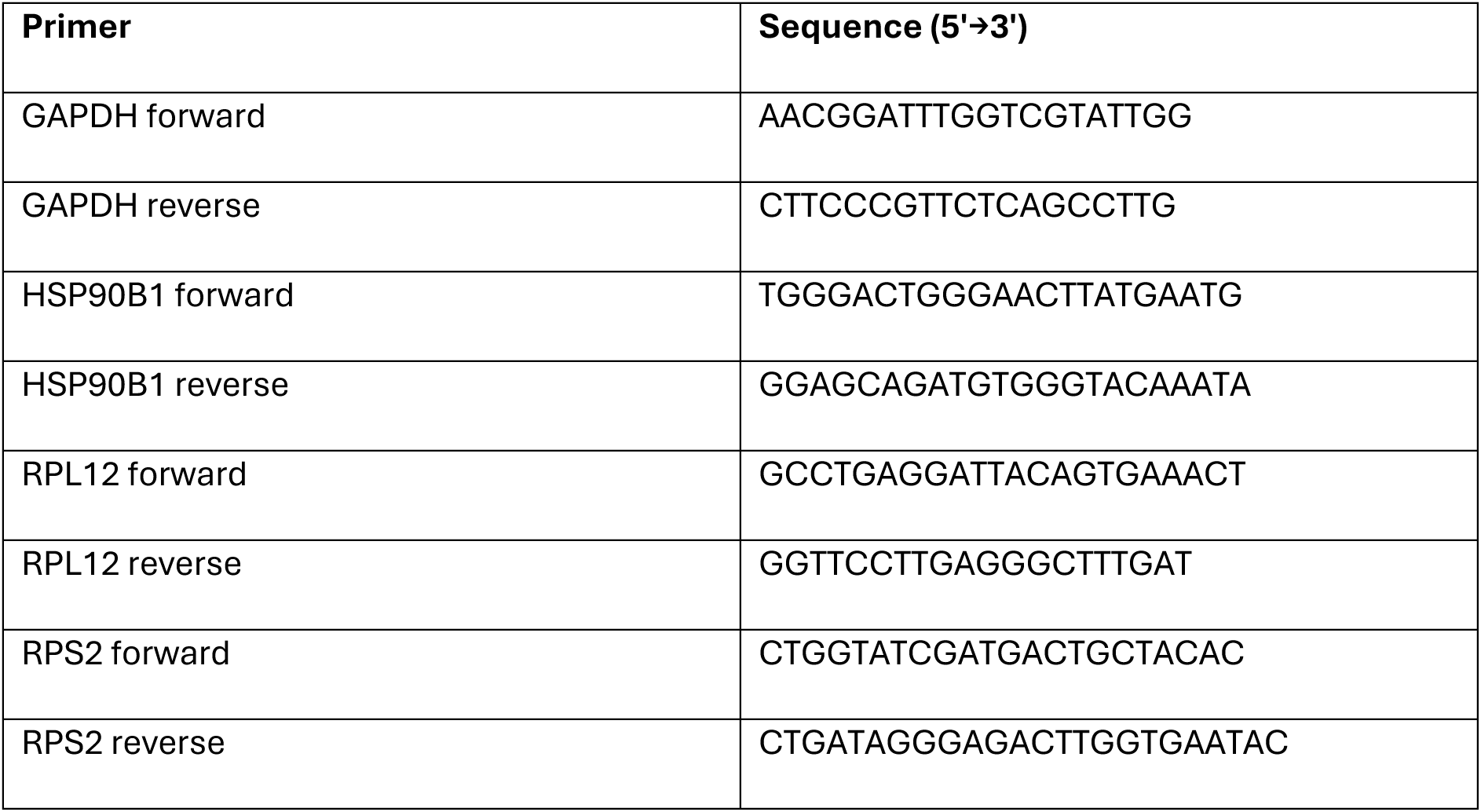
List of primers used in this study.

### Immunofluorescence

For immunofluorescence, cells were either grown and stained in LabTek chambers (Sigma-Aldrich, C7182) or cultured on glass coverslips in 24-well plates. Cells were briefly washed with cold PBS without calcium and magnesium and fixed in 4% formaldehyde for 15 min at room temperature, followed by three washes in 1× PBS. Permeabilization and blocking were performed for 1 h at room temperature in 1× PBS containing 0.5% Triton X-100 and 5% FBS. Primary antibodies were diluted in the same buffer and incubated overnight at 4°C, then washed off with three 10-min washes in PBS. Secondary antibodies were diluted 1:1000 in blocking buffer and applied for 1 h at room temperature, followed by three additional 10-min PBS washes. Slides were mounted in Duolink in situ mounting medium with DAPI (Sigma-Aldrich, DUO82020). Images were acquired on a Zeiss Axio Imager.M2 microscope (490020-0004-000) and analysed using the open-source ImageJ software (Fiji).

The following primary antibodies were used: mouse anti-Oct4 (STEMCELL Technologies, 60093; 1:1000), rabbit anti-Pax6 (STEMCELL Technologies, 60094; 1:500), rabbit anti-Calnexin (Cell Signaling Technology, 2679S; 1:50), and mouse anti-Tuj1 (STEMCELL Technologies, 60092; 1:500).

## Acknowledgments

We thank Frederic Schrøder Wenzel Arendrup for support with the RMS data analysis and Sophia J. Häfner for guidance on neuronal differentiations. Valuable advice on ER fractionation was received from Sarah L. Gillen and Chiara Giacomelli (Martin Bushell laboratory), who kindly shared a detailed version of their protocol. Henrik Nielsen also carefully read the manuscript and provided valuable comments, for which we are grateful.

This work was supported by the European Union’s Horizon 2020 research and innovation programme under the Marie Skłodowska-Curie grant agreement No. 945322 (LEAD) and by the Lundbeck Foundation (R380-2021-977) to ÜU. We thank the Henrik Nielsen lab for providing the mutated *A. thaliana* tRNA ligase and access to the sequencing facility, which is funded by grants from the Carlsberg Foundation and the NEYE-Foundation to HN.

## Figure legends

**Supplemental Figure 1. Quantification of cytosolic versus ER-associated rRNA across cell types.**

Percentage contribution of cytosolic and ER fractions to total RNA absorbance (A260) in HEK293 cells, human NPCs, and mature neurons. Across all three cell types, the ER fraction consistently accounts for approximately 22% of total rRNA content, with values ranging from ∼21–23%. Bar graphs represent means of three biological replicates.

**Supplemental Figure 2. Reproducibility of RiboMeth-seq methylation measurements across biological replicates.**

(A–B) Pairwise comparisons of cytosolic and ER RiboMeth-seq (RMS) scores for three independent biological replicates of HEK293 cells.

(C-D) Equivalent replicate comparisons for NPCs show similarly high reproducibility in both compartments.

(E-F) Equivalent replicate comparisons for neurons show similarly high reproducibility in both compartments.

**Supplemental Figure 3. NPCs exhibit a modest compartment-specific difference in 18S:462 methylation.**

RMS quantification of 18S:462 methylation levels in NPC cytosolic and ER-associated ribosomes. Methylation levels decrease from 98% in cytosolic ribosomes to 83% in ER-associated ribosomes, representing the only significant compartment-specific difference detected in NPCs (p = 0.049).

**Supplemental Figure 4. Neurons display a compartment-specific methylation shift at 28S:2043.**

(A) RiboMeth-seq quantification of 28S:2043 methylation in neuronal cytosolic and ER-associated ribosomes. Methylation at this site is elevated in ER ribosomes (19%) compared with cytosolic ribosomes (∼3%), representing the only significant neuron-specific compartment difference (p = 0.009).

(B) Normalized 5′- and 3′-end read coverage in a ±10 nt window surrounding 28S:2043. Cytosolic and ER-associated fractions are overlaid; for each read series, signals in both compartments are normalized across the window so that relative coverage differences between compartments are preserved. The dashed vertical line marks nucleotide 2043. For 28S rRNA numbering, see Materials and Methods and Supplemental Table 1.

## References

Aspden, Julie, William James Faller, Maria Barna, and Anders Lund. 2025. “Ribosome Heterogeneity and Specialization.” Philosophical Transactions of the Royal Society B: Biological Sciences 380(1921):20230375. doi:10.1098/rstb.2023.0375.

Bergeron, Danny, Hermes Paraqindes, Étienne Fafard-Couture, Gabrielle Deschamps-Francoeur, Laurence Faucher-Giguère, Philia Bouchard-Bourelle, Sherif Abou Elela, Frédéric Catez, Virginie Marcel, and Michelle S. Scott. 2023. “snoDB 2.0: An Enhanced Interactive Database, Specializing in Human snoRNAs.” Nucleic Acids Research 51(D1):D291–96. doi:10.1093/nar/gkac835.

Birkedal, Ulf, Mikkel Christensen-Dalsgaard, Nicolai Krogh, Radhakrishnan Sabarinathan, Jan Gorodkin, and Henrik Nielsen. 2015. “Profiling of Ribose Methylations in RNA by High-Throughput Sequencing.” Angewandte Chemie International Edition 54(2):451–55. doi:10.1002/anie.201408362.

Blair, John D., Dirk Hockemeyer, Jennifer A. Doudna, Helen S. Bateup, and Stephen N. Floor. 2017. “Widespread Translational Remodeling during Human Neuronal Differentiation.” Cell Reports 21(7):2005–16. doi:10.1016/j.celrep.2017.10.095.

Bourke, Ashley M., Andre Schwarz, and Erin M. Schuman. 2023. “De-Centralizing the Central Dogma: mRNA Translation in Space and Time.” Molecular Cell 83(3):452–68. doi:10.1016/j.molcel.2022.12.030.

Braakman, I., and D. N. Hebert. 2013. “Protein Folding in the Endoplasmic Reticulum.” Cold Spring Harbor Perspectives in Biology 5(5):a013201–a013201. doi:10.1101/cshperspect.a013201.

Chen, Qiang, Sujatha Jagannathan, David W. Reid, Tianli Zheng, and Christopher V. Nicchitta. 2011. “Hierarchical Regulation of mRNA Partitioning between the Cytoplasm and the Endoplasmic Reticulum of Mammalian Cells” edited by R. Hegde. Molecular Biology of the Cell 22(14):2646–58. doi:10.1091/mbc.e11-03-0239.

Child, Jessica R., Qiang Chen, David W. Reid, Sujatha Jagannathan, and Christopher V. Nicchitta. 2021. “Recruitment of Endoplasmic Reticulum-Targeted and Cytosolic mRNAs into Membrane-Associated Stress Granules.” RNA 27(10):1241–56. doi:10.1261/rna.078858.121.

Christopher, Josie A., Charlotte Stadler, Claire E. Martin, Marcel Morgenstern, Yanbo Pan, Cora N. Betsinger, David G. Rattray, Diana Mahdessian, Anne-Claude Gingras, Bettina Warscheid, Janne Lehtiö, Ileana M. Cristea, Leonard J. Foster, Andrew Emili, and Kathryn S. Lilley. 2021. “Subcellular Proteomics.” Nature Reviews Methods Primers 1(1):32. doi:10.1038/s43586-021-00029-y.

Gay, David M., Anders H. Lund, and Martin D. Jansson. 2022. “Translational Control through Ribosome Heterogeneity and Functional Specialization.” Trends in Biochemical Sciences 47(1):66–81. doi:10.1016/j.tibs.2021.07.001.

Gemmer, Max, Marten L. Chaillet, Joyce Van Loenhout, Rodrigo Cuevas Arenas, Dimitrios Vismpas, Mariska Gröllers-Mulderij, Fujiet A. Koh, Pascal Albanese, Richard A. Scheltema, Stuart C. Howes, Abhay Kotecha, Juliette Fedry, and Friedrich Förster. 2023. “Visualization of Translation and Protein Biogenesis at the ER Membrane.” Nature 614(7946):160–67. doi:10.1038/s41586-022-05638-5.

Genuth, Naomi R., and Maria Barna. 2018. “The Discovery of Ribosome Heterogeneity and Its Implications for Gene Regulation and Organismal Life.” Molecular Cell 71(3):364–74. doi:10.1016/j.molcel.2018.07.018.

Häfner, Sophia J., Martin D. Jansson, Kübra Altinel, Kasper L. Andersen, Zehra Abay-Nørgaard, Patrice Ménard, Martin Fontenas, Daniel M. Sørensen, David M. Gay, Frederic S. Arendrup, Disa Tehler, Nicolai Krogh, Henrik Nielsen, Matthew L. Kraushar, Agnete Kirkeby, and Anders H. Lund. 2023. “Ribosomal RNA 2′-O-Methylation Dynamics Impact Cell Fate Decisions.” Developmental Cell 58(17):1593–1609.e9. doi:10.1016/j.devcel.2023.06.007.

Holt, Christine E., Kelsey C. Martin, and Erin M. Schuman. 2019. “Local Translation in Neurons: Visualization and Function.” Nature Structural & Molecular Biology 26(7):557–66. doi:10.1038/s41594-019-0263-5.

Horste, Ellen L., Mervin M. Fansler, Ting Cai, Xiuzhen Chen, Sibylle Mitschka, Gang Zhen, Flora C. Y. Lee, Jernej Ule, and Christine Mayr. 2023. “Subcytoplasmic Location of Translation Controls Protein Output.” Molecular Cell 83(24):4509–4523.e11. doi:10.1016/j.molcel.2023.11.025.

Jansson, Martin D., Sophia J. Häfner, Kübra Altinel, Disa Tehler, Nicolai Krogh, Emil Jakobsen, Jens V. Andersen, Kasper L. Andersen, Erwin M. Schoof, Patrice Ménard, Henrik Nielsen, and Anders H. Lund. 2021. “Regulation of Translation by Site-Specific Ribosomal RNA Methylation.” Nature Structural & Molecular Biology 28(11):889–99. doi:10.1038/s41594-021-00669-4.

Krogh, Nicolai, Fazila Asmar, Christophe Côme, Helga Fibiger Munch-Petersen, Kirsten Grønbæk, and Henrik Nielsen. 2020. “Profiling of Ribose Methylations in Ribosomal RNA from Diffuse Large B-Cell Lymphoma Patients for Evaluation of Ribosomes as Drug Targets.” NAR Cancer 2(4):zcaa035. doi:10.1093/narcan/zcaa035.

Krogh, Nicolai, Martin D. Jansson, Sophia J. Häfner, Disa Tehler, Ulf Birkedal, Mikkel Christensen-Dalsgaard, Anders H. Lund, and Henrik Nielsen. 2016. “Profiling of 2′- *O* -Me in Human rRNA Reveals a Subset of Fractionally Modified Positions and Provides Evidence for Ribosome Heterogeneity.” Nucleic Acids Research 44(16):7884–95. doi:10.1093/nar/gkw482.

Luo, Jingchuan, Stuti Khandwala, Jingjie Hu, Song-Yi Lee, Kelsey L. Hickey, Zebulon G. Levine, J. Wade Harper, Alice Y. Ting, and Jonathan S. Weissman. 2025. “Proximity-Specific Ribosome Profiling Reveals the Logic of Localized Mitochondrial Translation.” Cell 188(20):5589–5604.e17. doi:10.1016/j.cell.2025.08.002.

Marcel, Virginie, Janice Kielbassa, Virginie Marchand, Kundhavai S. Natchiar, Hermes Paraqindes, Flora Nguyen Van Long, Lilia Ayadi, Valérie Bourguignon-Igel, Piero Lo Monaco, Déborah Monchiet, Véronique Scott, Laurie Tonon, Susan E. Bray, Alexandra Diot, Lee B. Jordan,Alastair M. Thompson, Jean-Christophe Bourdon, Thierry Dubois, Fabrice André, Frédéric Catez, Alain Puisieux, Yuri Motorin, Bruno P. Klaholz, Alain Viari, and Jean-Jacques Diaz. 2020. “Ribosomal RNA 2′O-Methylation as a Novel Layer of Inter-Tumour Heterogeneity in Breast Cancer.” NAR Cancer 2(4):zcaa036. doi:10.1093/narcan/zcaa036.

Morin, Chloé, Hermes Paraqindes, Flora Nguyen Van Long, Caroline Isaac, Emilie Thomas, Dennis Pedri, Carlos Ariel Pulido-Vicuna, Anne-Pierre Morel, Virginie Marchand, Yuri Motorin, Marjorie Carrere, Jessie Auclair, Valéry Attignon, Roxane M. Pommier, Emmanuelle Ruiz, Fleur Bourdelais, Frédéric Catez, Sébastien Durand, Anthony Ferrari, Alain Viari, Jean-Christophe Marine, Alain Puisieux, Jean-Jacques Diaz, Caroline Moyret-Lalle, and Virginie Marcel. 2025. “Specific Modulation of 28S_Um2402 rRNA 2′- *O* -Ribose Methylation as a Novel Epitranscriptomic Marker of ZEB1-Induced Epithelial–Mesenchymal Transition in Different Mammary Cell Contexts.” NAR Cancer 7(1):zcaf001. doi:10.1093/narcan/zcaf001.

Reid, David W., and Christopher V. Nicchitta. 2012. “Primary Role for Endoplasmic Reticulum-Bound Ribosomes in Cellular Translation Identified by Ribosome Profiling.” Journal of Biological Chemistry 287(8):5518–27. doi:10.1074/jbc.M111.312280.

Reid, David W., and Christopher V. Nicchitta. 2015. “Diversity and Selectivity in mRNA Translation on the Endoplasmic Reticulum.” Nature Reviews Molecular Cell Biology 16(4):221–31. doi:10.1038/nrm3958.

Sánchez, Wendy N., Arnold J. M. Driessen, and Christian A. M. Wilson. 2025. “Protein Targeting to the ER Membrane: Multiple Pathways and Shared Machinery.” Critical Reviews in Biochemistry and Molecular Biology 1–47. doi:10.1080/10409238.2025.2503746.

Sloan, Katherine E., Ahmed S. Warda, Sunny Sharma, Karl-Dieter Entian, Denis L. J. Lafontaine, and Markus T. Bohnsack. 2017. “Tuning the Ribosome: The Influence of rRNA Modification on Eukaryotic Ribosome Biogenesis and Function.” RNA Biology 14(9):1138–52. doi:10.1080/15476286.2016.1259781.

Stephens, Samuel B., and Christopher V. Nicchitta. 2008. “Divergent Regulation of Protein Synthesis in the Cytosol and Endoplasmic Reticulum Compartments of Mammalian Cells” edited by M. P. Wickens. Molecular Biology of the Cell 19(2):623–32. doi:10.1091/mbc.e07-07-0677.

Yang, Jiabin, Zhongyang Chen, Jiayin He, Binbin Zou, Yanmin Si, Yanni Ma, and Jia Yu. 2025. “Modular RNA Interactions Shape FXR1 Condensates Involved in mRNA Localization and Translation.” Nature Communications 16(1):8589. doi:10.1038/s41467-025-63700-y.

Zhou, Fengbiao, Nesrine Aroua, Yi Liu, Christian Rohde, Jingdong Cheng, Anna-Katharina Wirth, Daria Fijalkowska, Stefanie Göllner, Michelle Lotze, Haiyang Yun, Xiaobing Yu, Caroline Pabst, Tim Sauer, Thomas Oellerich, Hubert Serve, Christoph Röllig, Martin Bornhäuser, Christian Thiede, Claudia Baldus, Michaela Frye, Simon Raffel, Jeroen Krijgsveld, Irmela Jeremias, Roland Beckmann, Andreas Trumpp, and Carsten Müller-Tidow. 2023. “A Dynamic rRNA Ribomethylome Drives Stemness in Acute Myeloid Leukemia.” Cancer Discovery 13(2):332–47. doi:10.1158/2159-8290.CD-22-0210.

Zinnall, Ulrike, Miha Milek, Igor Minia, Carlos H. Vieira-Vieira, Simon Müller, Guido Mastrobuoni, Orsalia-Georgia Hazapis, Simone Del Giudice, David Schwefel, Nadine Bley, Franka Voigt, Jeffrey A. Chao, Stefan Kempa, Stefan Hüttelmaier, Matthias Selbach, and Markus Landthaler. 2022. “HDLBP Binds ER-Targeted mRNAs by Multivalent Interactions to Promote Protein Synthesis of Transmembrane and Secreted Proteins.” Nature Communications 13(1):2727. doi:10.1038/s41467-022-30322-7.

